# Effects of BMP-2 on neovascularization during large bone defect regeneration

**DOI:** 10.1101/464396

**Authors:** HB Pearson, DE Mason, CD Kegelman, L Zhao, JH Dawahare, MA Kacena, JD Boerckel

## Abstract

Insufficient blood vessel supply is a primary limiting factor for regenerative approaches to large bone defect repair. Recombinant BMP-2 delivery induces robust bone formation and has been observed to enhance neovascularization, but whether the angiogenic effects of BMP-2 are due to direct endothelial cell stimulation or to indirect paracrine signaling remains unclear. Here, we evaluated the effects of BMP-2 delivery on vascularized bone regeneration and tested whether BMP-2 induces neovascularization directly or indirectly. We found that delivery of BMP-2 (5 μg) enhanced both bone formation and neovascularization in critically sized (8 mm) rat femoral bone defects; however, BMP-2 did not directly stimulate angiogenesis *in vitro*. In contrast, conditioned medium from both mesenchymal progenitor cells and osteoblasts induced angiogenesis *in vitro*, suggesting a paracrine mechanism of BMP-2 action. Consistent with this inference, co-delivery of BMP-2 with endothelial colony forming cells (ECFCs) to a heterotopic site, distant from the bone marrow niche, induced ossification but had no effect on neovascularization. Taken together, these data suggest that BMP-2 induces neovascularization during bone regeneration primarily through paracrine activation of osteoprogenitor cells.

## Introduction

Critical-sized bone defects that do not heal without intervention have prolonged morbidity and are extremely costly.^1,2^ Approximately 1.5 million bone grafting operations are performed annually in the United States.^3^ Although autologous bone is the ‘gold standard’ for bone grafting in segmental bone loss and intervertebral fusion, this treatment is limited by insufficient supply for large defects as well as substantial donor site pain and morbidity.^4^ Allografts are therefore often required to bridge the defect, but with limitations including graft rejection, disease transmission, and lower rates of healing compared to autografts due to a failure to revascularize and remodel, leading to re-fracture.^5^ Alternate regenerative strategies are needed, and tissue engineering has emerged as a promising alternative.

The most successful bone tissue engineering strategy to date is biomaterial-based delivery of osteoinductive growth factors including members of the bone morphogenetic protein family, such as BMP-2. BMP-2 is clinically approved for lumbar spine fusion, open tibial fractures and some non-unions, and maxillofacial regeneration, but is often used off-label for large segmental defects and fractures in patients with co-morbidities likely to produce non-union.^6,7^ This approach, in which soluble growth factor is delivered via absorbable collagen sponge, robustly induces new bone formation through osteoprogenitor cell recruitment and local stimulation of osteogenic differentiation and matrix production.^8,9^

New blood vessel formation and volumetric perfusion is critical for the regeneration of large bone defects as osteoblasts require < 300 μm proximity to a capillary supply for survival.^10^ BMP-2 delivery enhances new blood vessel formation in bone defects,^11^ but whether BMP-2 directly stimulates endothelial cell angiogenesis or indirectly through paracrine signaling from activated osteoprogenitor cells remains controversial.^12–16^

In this study, we evaluated the effects of BMP-2 delivery on neovascularization of large bone defects and whether BMP-2 directly induces endothelial cell activation or indirectly activates paracrine signaling from osteogenic cells. Here, we show that delivery of BMP-2 (5 μg per defect), enhanced both bone formation and neovascularization in large femoral bone defects; however, BMP-2 did not directly stimulate angiogenesis *in vitro*. In contrast, human MSC and mouse osteoblast-conditioned medium induced endothelial cell migration and tubular network formation *in vitro*, suggesting a paracrine mechanism. Finally, BMP-2 co-delivery with endothelial colony forming cells (ECFCs) to a heterotopic site, distant from the bone marrow niche, induced ossification but had no effect on neovascularization. Together these data suggest that BMP-2-induced neovascularization during bone regeneration occurs secondary to mesenchymal progenitor cell activation and bone formation rather than direct endothelial cell activation.

## Materials and Methods

### Rat femoral segmental bone defect model

Critical-sized (8mm) bilateral femoral defects were created in 14-week-old female SASCO Sprague Dawley rats and stabilized with custom internal fixation plates as described previously.^17^ Briefly, fixation plates were affixed to the anterolateral surface using four bi-cortical screws. Mid-diaphyseal segmental defects, 8 mm in length, were created using an oscillating saw under saline irrigation. Lyophilized Type I collagen scaffolds (average pore size 61.7 μm, 93.7% pore volume; DSM Biomedical) were cut into cylinders (h=9mm, d=5mm) and hydrated with 100 μl of phosphate-buffered saline (PBS) containing BMP-2, or rat serum albumin carrier. BMP-2 was purchased (R&D Systems, Medtronic), reconstituted in sterile PBS and diluted to 5 μg/100 μL. The hydrated collagen scaffolds were then placed in the femoral defect and the muscles and incision were sutured closed and secured with wound clips. Wound clips were removed 2 weeks post-surgery. For pain management, animals were given subcutaneous injections of buprenorphine (0.01-0.03 mg/kg) every 8 h for three days. All procedures were reviewed and approved by the Institutional Animal Care and Use Committees (IACUC), which follows the National Institutes of Health guidelines on the human care and use of laboratory animals.

### *In vivo* microCT imaging

Longitudinal microCT imaging was performed at 4 and 8 weeks post-surgery. Scanning was performed at high resolution, using voxel sizes of either 20 or 39 μm (depending on experiment), 79.9 mm FOV diameter 70 kVp 114 μA, and 8 W energy. Measurements were calibrated at 70 kVp, BH 1200 mgHA/ccm scaling with a 200 ms integration time for a slice thickness of 0.02 or 0.039 mm.

The defect region of interest was defined by the minimum distance across all samples using contours that captured all bone formed in each transverse slice. Bone formation at the proximal and distal ends of non-bridged samples were also assessed for bone volume and bone mineral density by individual contouring, beginning at the last slice containing original cortical bone and continuing to the last slice containing new bone forming the capped end.

### MicroCT angiography

Contrast-enhanced microCT angiography was performed to quantify and visualize three-dimensional (3D) neovascular structures, as described previously.^11^ Briefly, the ascending aorta was catheterized through the left ventricle of the heart, and serial solutions of heparin (200 ml), 10% neutral buffered formalin (100 ml) and PBS (100 ml). The vasculature was then manually perfused with the lead chromate-containing radiodense contrast agent, Microfil MV-122 (Flowtech). The contrast agent was diluted (4:1) to attenuate X-rays at the same threshold as newly-formed bone, to enable simultaneous reconstruction.^11,18^

MicroCT scans were performed to register both bone and vascular structures together using a voxel size of 15.6 or 20 μm, medium resolution, 31.9 mm FOV diameter, 70 kvp 114 μA, and 8 W energy. Measurements were calibrated at 70 kVp, BH 1200 mgHA/ccm scaling with a 200 ms integration time for a slice thickness of 0.0156 or 0.02 mm. Next, the mineralized tissues were fully decalcified using EDTA or Cal-Ex I (Fisher Scientific) with gentle agitation at room temperature for 4-8 weeks. The limbs were then re-scanned to image and quantify the vasculature alone. Bone volume in each defect was then calculated by subtracting the vessels-only scan from the composite bone-and-vessel scan.

The bone volume was analyzed using a 10mm diameter contour circles to ensure inclusion of all bone formed. The vessel analysis was performed using a 5mm diameter contour circle centered in the defect.

### *In vivo* experimental design

Three separate *in vivo* experiments were performed. The first two featured BMP-2 delivery to femoral segmental defects, and the third evaluated vascularized bone formation in a heterotopic (subcutaneous) site.

### Segmental defect study 1

Bilateral segmental bone defects were created in the femora of eight rats. Each animal received a treatment condition in one limb (5 μg BMP-2) and PBS control in the contralateral limb. MicroCT angiography was performed at week 3 post-surgery. One animal was euthanized due to complications during surgery, and two animals were excluded from vascular analysis due to incomplete contrast agent perfusion. One PBS-treated limb was also excluded from vascular analysis due to ruptured vessels during perfusion. Vessel analysis and statistics were therefore completed on five animals (N=4-5).

### Segmental defect study 2

Bilateral segmental bone defects were created in the femora of sixteen rats. Six animals received PBS-only in both limbs to assess potential systemic effects of BMP-2 treatment. Ten animals received BMP-2 treatment condition in one limb (5 μg BMP-2) and PBS control in the contralateral limb (N = 10 for BMP-2/PBS paired). In this study, *in vivo* microCT analysis was performed at weeks 4 and 8 post-surgery, and high-resolution microCT angiography was then performed at week 8 post-surgery. Four animals died due to complications, and three animals were excluded from microCT angiography analysis due to incomplete contrast agent perfusion. Longitudinal microCT analysis and statistics were evaluated in 12 animals (BMP-2 treated N = 9, PBS treated N = 3). Vessel analysis and statistics were evaluated in 9 animals (BMP-2 treated N=7, PBS-only treated N = 2).

### Subcutaneous study

Type I collagen hydrogel matrices (2.5 mg/ml, GeniPhys) were polymerized in 24 well-plates into disc-shaped constructs of 240μl volume. Matrices were polymerized containing four conditions: with or without endothelial colony forming cells (ECFCs; 1.6×10^6^ cells/ml; 384,000 cells per construct) and with or without BMP-2 (100 ng/ml; 24 ng per construct). Collagen matrices were then cultured for 24 hours prior to implantation in 500 μl endothelial growth medium (EGM-2) containing 10% fetal bovine serum (FBS). For implantation of human ECFCs in vivo, Rowett Nude (RNU) rats were used to minimize immune reactions. Fourteen-week-old female RNU rats (N = 6) received four implants per animal, with two implants placed subcutaneously over each back flank, respectively. Briefly, a single incision was made on the dorsum, and two separate pockets were blunt dissected above each flank such that each subcutaneous pocket was separated by intact fascia. The hydrogels were placed inside custom polysulfone rings to enable region of interest determination (h=1.77 di=11mm, do=13mm) and deposited into each pocket. Conditions were evenly randomized between animals. For pain management, animals were given subcutaneous injections of buprenorphine (0.01 mg/kg) every 8 h for two days. MicroCT angiography was performed at week 4.

### ECFC wound migration assay

ECFC migration was evaluated in the wound migration assay, as described previously.^18,19^ Briefly, passage P6-P8 endothelial colony forming cells (ECFCs) were plated in 6 well plates and allowed to grow to 80% confluence. After a 2-hour serum starve in endothelial basal medium (EBM-2), wounds were created in the cell monolayer by dragging a 200uL pipet tip across the well both vertically and horizontally to form a cross. Cells were then washed with EBM-2 to remove any dead cells and supplemented with either EBM-2 only (negative control), 2% serum in EBM-2, VEGF (100 ng/ml) in EBM-2, or BMP-2 (100 ng/ml and 200 ng/ml) in EBM-2. Cell migration was monitored by taking images at 0, 4, 8, and 12 hours. Percent wound closure was calculated by measuring the wound area at each time point and dividing by the initial wound area at the time of scratch. Wound closure rate was calculated by multiplying the percent wound closure by the initial scratch width and dividing by the migration time.

### ECFC tubular network formation assay

Tubulogenesis was evaluated using the matrigel tube formation assay as described previously.^18,19^ Briefly, reduced growth factor basement membrane matrix (Trevigen 3433-001-R1) was polymerized in a 96 well plates (60μl) at 37°C for 30 minutes. Passage P6-P8 ECFCs were plated on the polymerized basement membrane matrix at 50,000 cells/cm^2^ (16,190 cells/well). Treatment media was added according to seven groups: EBM-2 only (control), 2% serum in EBM-2, or EBM-2 supplemented with VEGF (100ng/ml), or BMP-2 (100ng/ml, 200ng/ml). Cells were incubated at 37°C and imaged at 4 and 8 hours. Total vessel length, average vessel length, number of nodes, number of branches, and number of branches per node were measured using ImageJ.^20^

### SMAD1/5/9 immunolocalization

ECFCs were cultured for 24 hours on glass coverslips with an initial seeding density of 50,000 cells/cm^2^ and serum starved for 2 hours. Cells were then treated with 10 ng/ml of BMP-2 for 0, 15, 30, 45, and 60 min. Cells were fixed with 37°C pre-warmed 3.7% formaldehyde, 5% sucrose for 20 min at room temperature, permeabilized and blocked with 0.3% triton-x and 5% goat serum for 1 hour at room temperature. Cells were washed with PBS and treated with SMAD 1/5/9 primary antibody at a 1:200 dilution (abcam #ab80255) at 4°C overnight and visualized secondary Alex Fluor 594 at 10 μg/ml for 30 min at room temperature. Cells were counterstained with FITC-phalloidin (to visualize F-actin) and DAPI (nuclei). SMAD intensity and localization was quantified using ImageJ.^20^

### Conditioned media experiments

Wound migration and tube formation experiments were repeated using conditioned media from bone marrow stromal cells (hMSC) and/or mouse calvarial osteoblasts (Ob). First, conditioned media was isolated from hMSCs (Lonza) plated in 6 well plates for 3 or 6 days with or without 100 ng/ml of BMP-2. EBM-2 was then supplemented at 30% for migration and tube formation assays. Next, mouse calvarial osteoblast-conditioned media experiments were also performed using media from 10cm plates 400k calvarial cells, cultured for 6 days. Mouse calvarial cells were isolated and cultured as described previously.^21,22^ Basal media was a-MEM with 10% serum and penicillin-streptomycin-glutamine. EBM-2 was supplemented with condition media at 10 or 30% for migration assays and at 30% for tube formation assays.

### Statistical methods

Differences between groups were evaluated by paired Student’s t-test or one way analysis of variance (ANOVA) followed by Tukey’s post-hoc test for multiple comparisons, when assumptions for homoscedasticity and normality were met by Brown-Forsythe and D’Agostino and Pearson tests, respectively. When assumptions for normality and homoscedasticity were not met, data was log transformed and the parametric analyses were repeated. Kruskal-Wallis non-parametric and Dunn’s multiple comparisons tests were used when normality and homoscedasticity conditions were not achieved by log transformation. P-values less than 0.05 were considered significant. Significance indicator letters shared in common between groups indicate no significant difference in pairwise comparisons.

## Results

The objectives of this study were first to determine the extent to which BMP-2 enhances vascularized bone regeneration and second to test whether the neovascular response to BMP-2 treatment is direct (by stimulating endothelial cell angiogenesis) or indirect (via paracrine signaling from other cells).

### Effects of BMP-2 on vascularized bone regeneration at week 3

In our first experiment, we delivered BMP-2 (5 μg) to large bone defects and evaluated bone formation and neovascularization at week 3 post-surgery. BMP-2 delivery significantly increased bone formation compared to contralateral PBS controls (**Fig. 1**).

**Figure 1.**
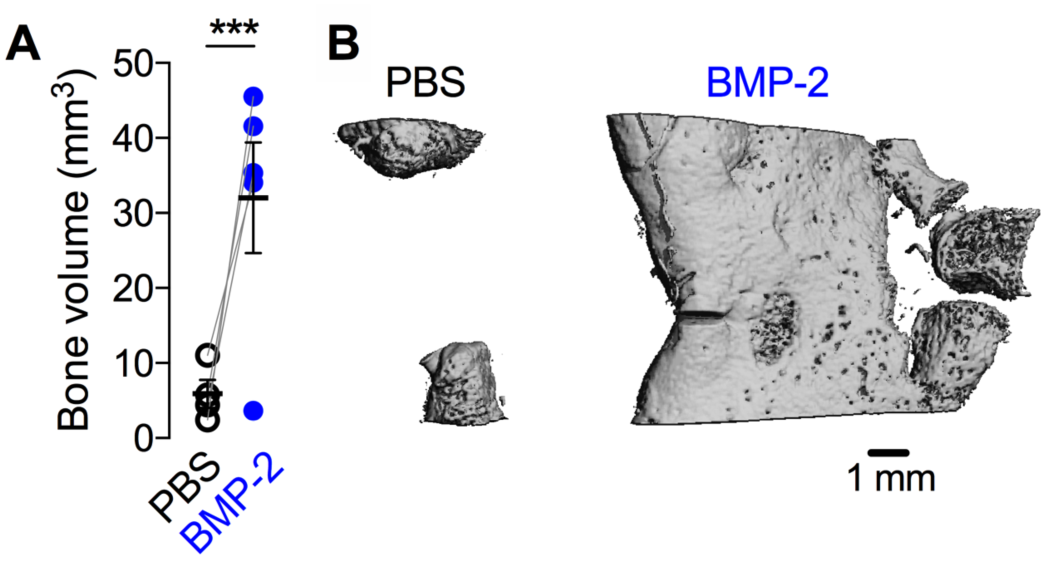
Effects of 5 μg BMP-2 delivery on bone formation at week 3. **A**) MicroCT-measured bone volume. Comparisons among and between groups were performed by two-way ANOVA with matched pairs and Bonnferroni’s multiple comparisons test. N = 4-5 per group. Each replicate is shown with summary statistics represented as mean ± s.e.m. *** indicates p ≤ 0.001 vs. all other groups. **B**) Representative microCT images of bone formation in the defect.

To determine whether BMP-2 enhanced blood vessel formation, we quantified the amount and morphology of the neovascular structures in the defect at week 3 by microCT angiography. BMP-2 delivery significantly increased neovascular volume fraction, vascular connectivity, and vessel number, but diameter, and spacing were not significantly different (**Fig. 2A-F**).

**Figure 2.**
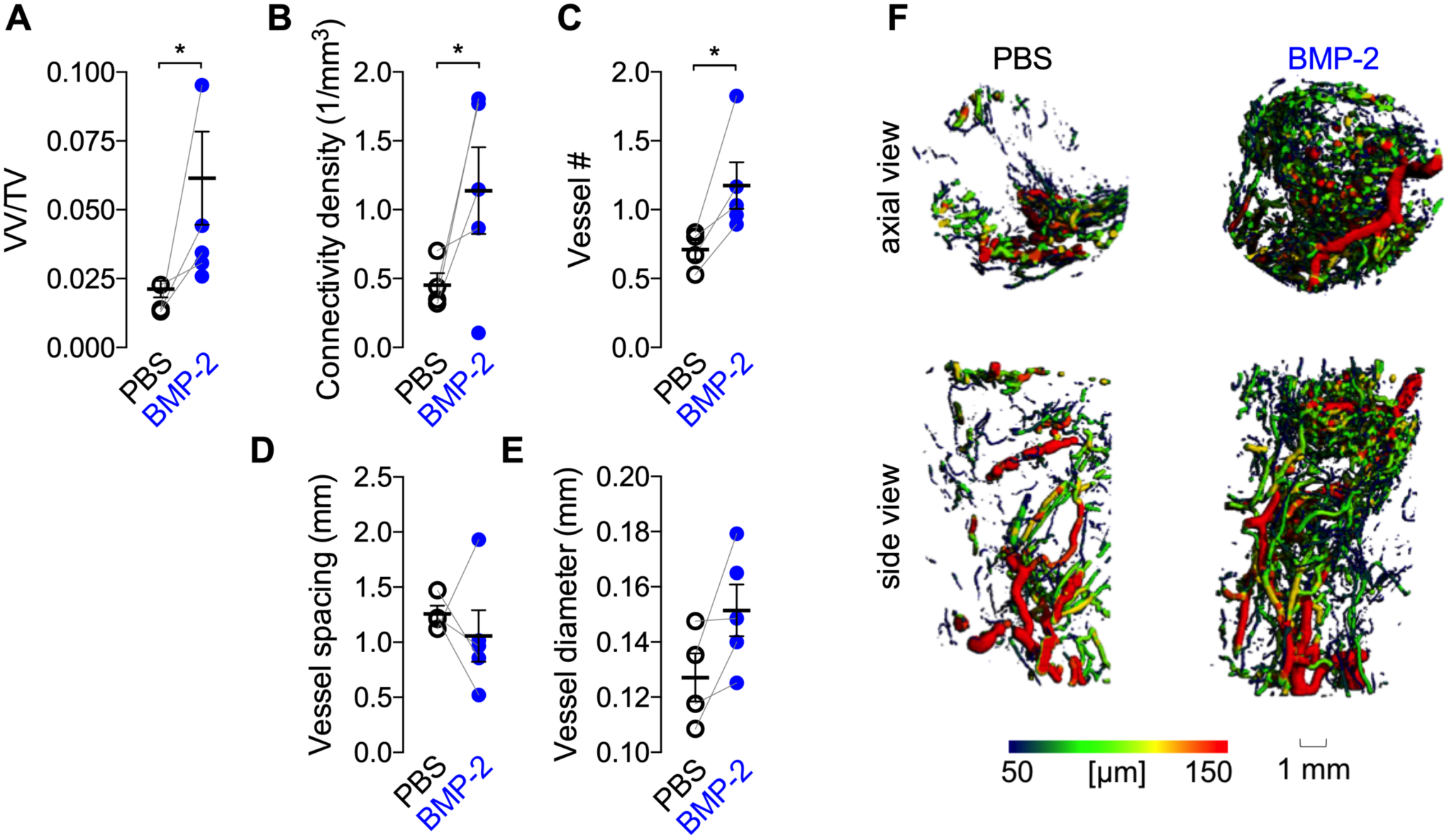
Effects of 5 μg BMP-2 delivery on blood vessel formation at week 3. **A**) Vessel volume fraction (vessel volume/total volume). **B**) Vascular connectivity density. **C**) Vessel number. **D**) Vessel spacing. **E**) Vessel diameter. Each replicate is shown with summary statistics represented as mean ± s.e.m. Paired limbs are connected by gray lines and mean and standard error of the mean are indicated in black. Comparisons among and between groups were performed by paired Student’s t-tests. N=4-5 per group. * indicates p≤0.05. **F**) Representative microCT images of defect vascularization with local vascular diameter mapping.

### Effects of BMP-2 on vascularized bone regeneration at week 8

We next evaluated the effects of BMP-2 (5 μg) delivery on vascularized bone regeneration over eight weeks by longitudinal *in vivo* microCT imaging and microCT angiography at week 8. BMP-2-treatment increased bone formation amount and rate compared to contralateral PBS controls (**Fig. 3A**). Animals with paired PBS-only scaffolds in both limbs showed similar bone formation to contralateral controls for BMP-2-treated limbs, suggesting no systemic effects of BMP-2 on bone regeneration. Post-mortem bone volume for BMP-2-treated limbs was significantly greater (p < 0.001) than PBS controls (**Fig. 3B**). Bone mineral density was significantly lower (p < 0.001) for the BMP-2 treated defects, indicative of new woven bone formation (**Fig. 3C**). Representative images of bone formation local mineral density mapping on a saggital cut-plate at week 8 are shown (**Fig 3D**).

**Figure 3.**
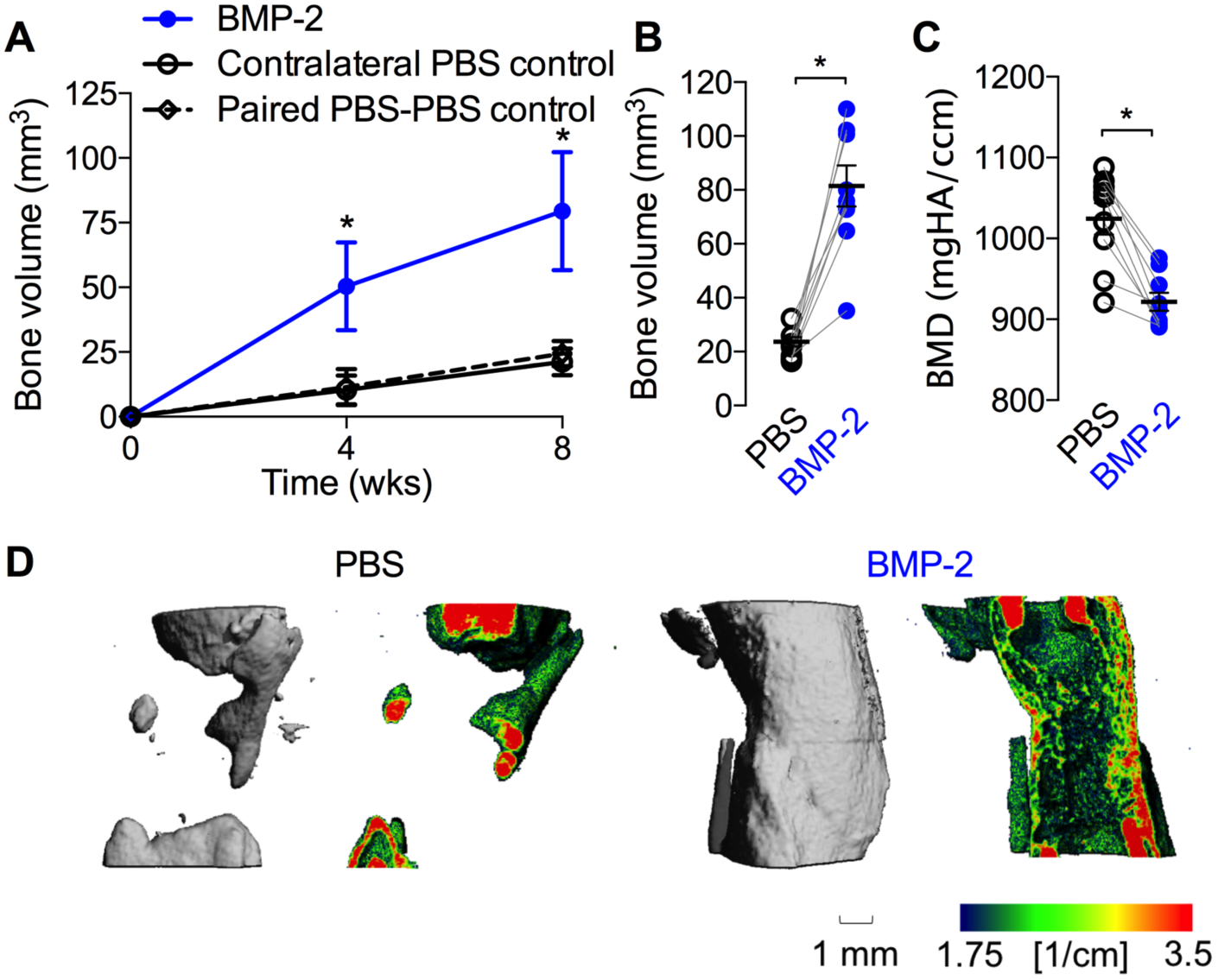
Effects of 5 μg BMP-2 delivery on bone formation at week 8. **A**) Longitudinal *in vivo* microCT-measured bone volume. Summary statistics represented as mean ± sem. **B**) Post-mortem bone volume at week 8. **C**) Post-mortem bone mineral density (BMD) at week 8. **D**) Representative microCT images of bone formation and local mineral density mapping. Each replicate is shown with summary statistics represented as mean ± s.e.m. Comparisons among and between groups were performed by repeated measures or two-way paired ANOVA and Bonnferroni’s multiple comparisons test. N = 9 per group, paired PBS-PBS controls, N=3. * indicates p ≤ 0.05 vs. all other groups at each time point. ***,**** indicate p < 0.001 and p < 0.0001, respectively.

By week 8, 5 μg BMP-2 treatment significantly elevated vascular volume fraction, connectivity, and number (**Fig. 4A-C**), and reduced vessel spacing and degree of anisotropy (**Fig. 4D,F**). Vessel diameter was not significantly different between groups (**Fig. 4E**). Animals with paired PBS only scaffolds in both limbs were not included in the quantitative analysis due to small sample size (N = 2), but their values were comparable to the contralateral controls of the BMP-2 animals (data not shown).

**Figure 4.**
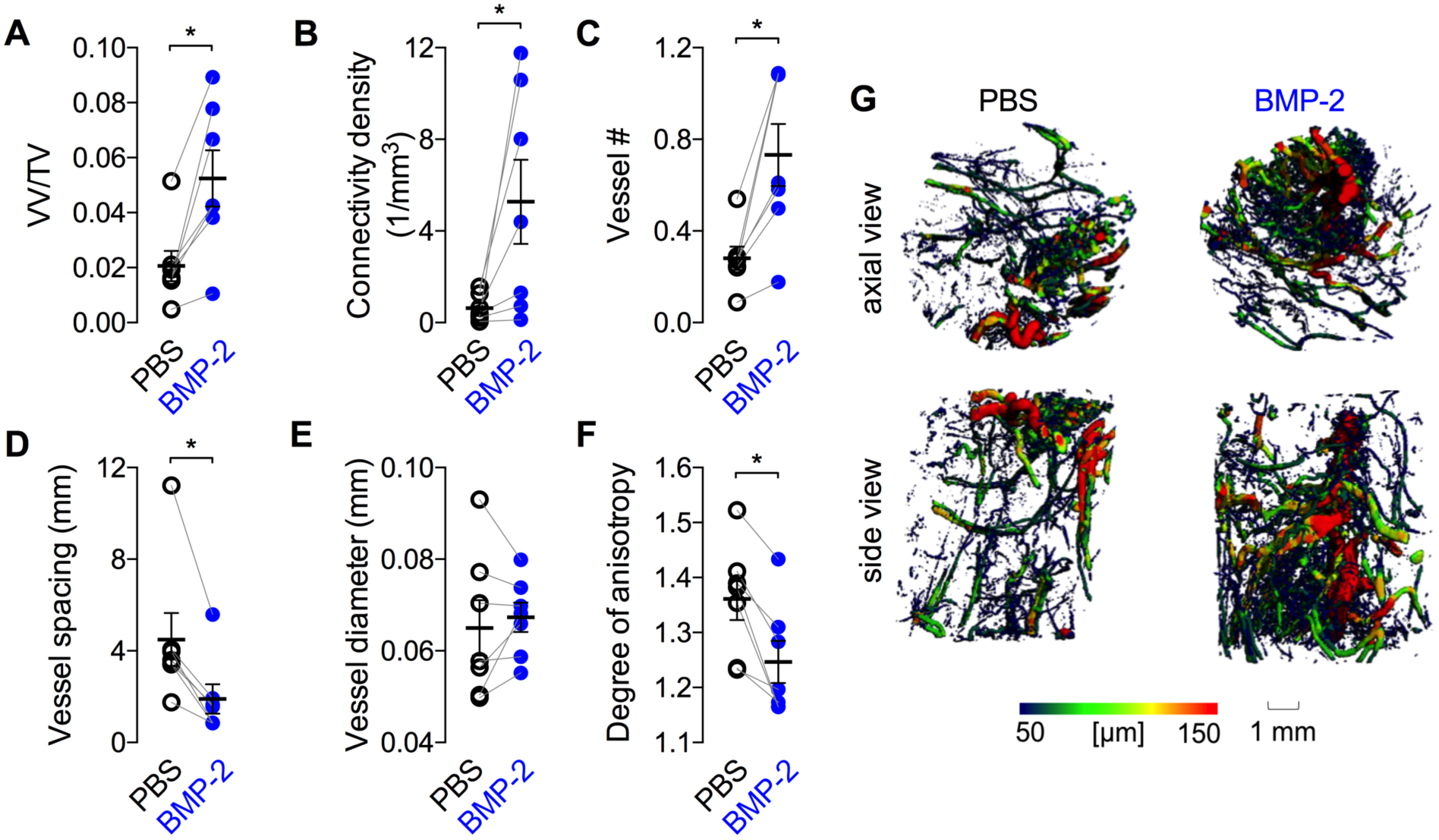
Effects of 5 μg BMP-2 delivery on blood vessel formation at 8 weeks. **A**) vessel volume fraction, **B**) connectivity density, **C**) vessel number, **D**) vessel diameter, **E**) vessel diameter, **F**) degree of anisotropy. Each replicate is shown with summary statistics represented as mean ± s.e.m. Paired limbs are connected by gray lines and mean values are indicated by the horizontal lines. *** indicates (p ≤ 0.001) and ** indicates (p < 0.01). Comparisons among and between groups were performed by paired Student’s t-tests. N=7-8 per group. **G**) Representative microCT images of defect vascularization with local vascular diameter mapping.

### ECFC responsiveness to BMP-2

BMP-2 signals, in part, by inducing nuclear localization of the transcription factors SMAD1/5/9. To test whether BMP-2 stimulation induces canonical SMAD signaling in endothelial cells, we quantified SMAD1/5/9 nuclear localization in ECFCs by immunofluorescence after 10 ng/ml of BMP-2 stimulation for 0, 15, 30, 45, and 60 minutes. BMP-2 treatment significantly increased the ratio of nuclear:cytosolic SMAD1/5/9 after 60 minutes (**Fig. 5**).

**Figure 5.**
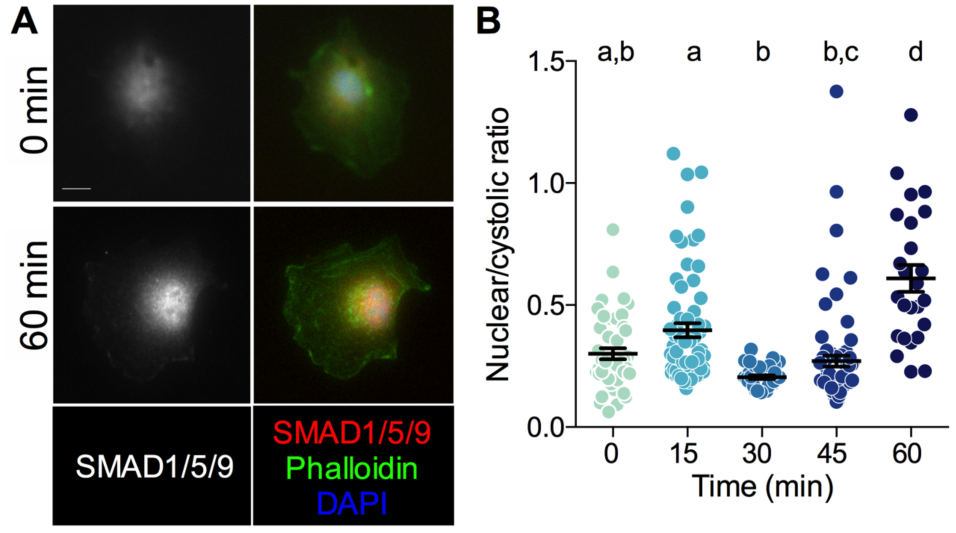
BMP-2 stimulation of SMAD1/5/9 nuclear localization in ECFCs. **A**) Representative images of SMAD1/5/9 staining at 0 and 60 min after BMP-2 treatment. **B**) Quantfication of nuclear/cytosolic ratio. Scale bar = 10 μm. Each replicate is shown with summary statistics represented as mean ± s.e.m. Significance indicator letters shared in common between groups indicate no significant difference in pairwise comparisons (p > 0.05) by Kruskal-Wallis and Dunn’s multiple comparisons test. N=25-76.

### Effects of BMP-2 on angiogenesis *in vitro*

We next tested whether BMP-2 induced signaling was effective to stimulate ECFC migration and tube formation *in vitro*. Cells were stimulated with either low (100 ng/ml) or high (200 ng/ml) doses of BMP-2 and cell migration and tube formation were quantified over 12 and 8 hours, respectively, in comparison to serum-free basal medium negative controls and VEGF (100 ng/ml) or 2% FBS positive controls. Both positive controls significantly induced migratory wound closure, but BMP-2 treatment did not alter cell migration (**Fig. 6A,B**). FBS positive controls significantly increased tube length, but BMP-2 treatment did not (**Fig. 6C-F**).

**Figure 6.**
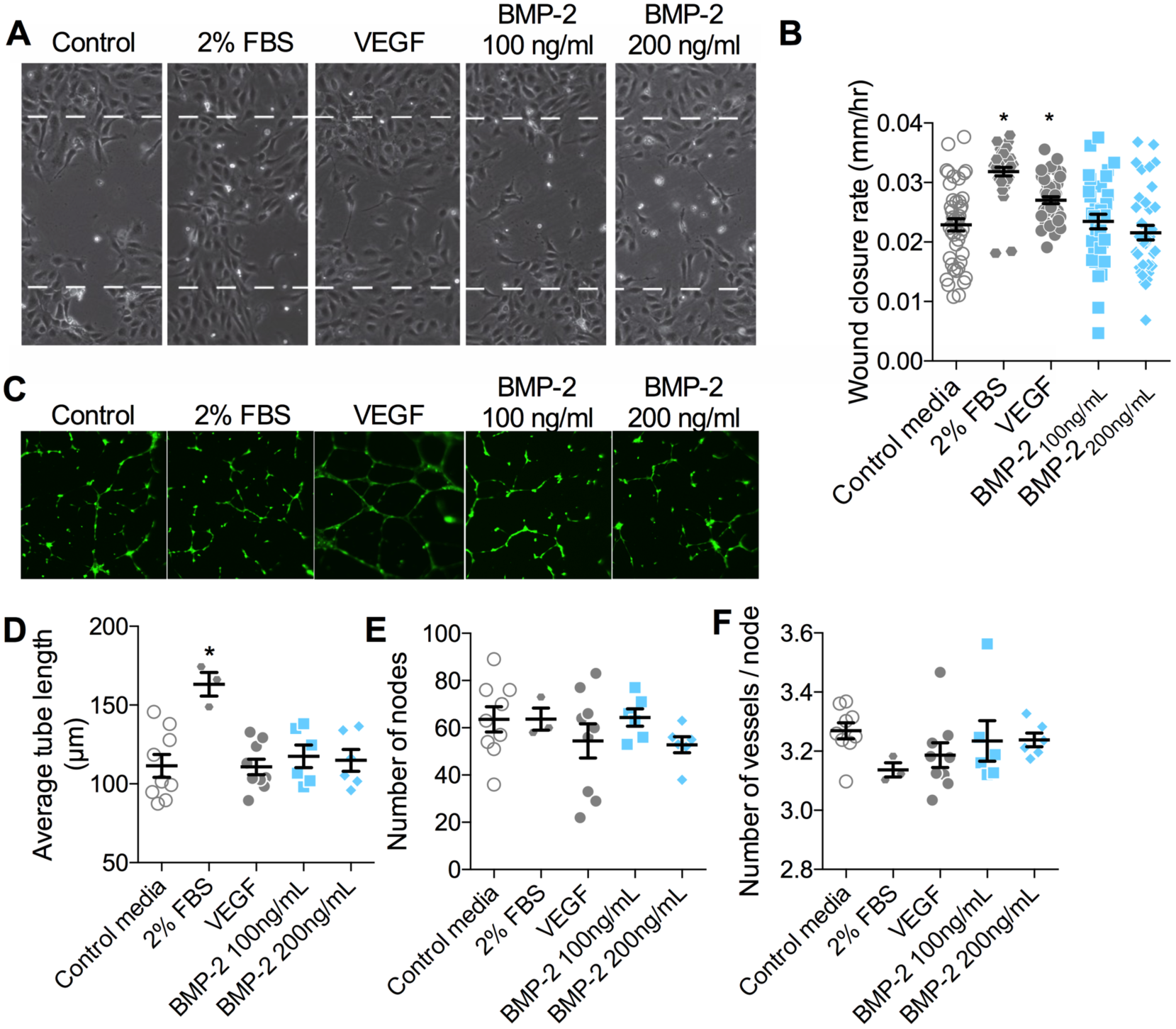
Effects of BMP-2 on endothelial cell migration and tube formation. A-B) ECFC wound closure was evaluated over 12 hours (**A**) and quantified as wound closure rate (**B**). All data points shown with mean and s.e.m. Initial scratch width indicated by dashed line (450 μm). * indicates p < 0.05 compared to negative control by Kruskal-Wallis and Dunn’s multiple comparisons test. n=36-44 replicates, in N = 6 independent experiments. **C-F**) ECFC tubulogenesis was evaluated over 8 hours and visualized by calcein-AM staining (**C**). Network formation was quantified by average tube length (**D**), number of nodes (**E**), and number of vessels per node (**F**). Each independent replicate is shown with summary statistics represented as mean ± s.e.m. Initial scratch width indicated by dashed line (450μm). * indicates p < 0.05 compared to negative control by ANOVA and Tukey’s multiple comparisons test. N=3-9.

### Effects of hMSC- and calvarial osteoblast-conditioned media on angiogenesis

Since BMP-2 stimulation failed to induce angiogenesis in isolated endothelial cells, we next tested the alternative hypothesis that BMP-2 induces angiogenesis by paracrine signaling from activated progenitor cells and osteoblasts. To this end, we cultured hMSCs with or without BMP-2 stimulation for 3 or 6 days and collected conditioned media, which we then mixed to 30% by volume with ECFC basal medium. We then treated ECFCs in the wound migration and tube formation assays with either hMSC-conditioned medium (hMSC-CM) or BMP-2-treated hMSC-conditioned medium (hMSC-CM + BMP-2) in comparison with 30% hMSC/70% ECFC basal medium (BM) as a control. Media conditioned for 3 days had no effect on ECFC migration (**Fig. 7A**) or tube formation (**Fig. 7B-D**), but media conditioned by hMSCs for 6 days significantly increased both ECFC migration speed and tubular network length (**Fig. 7A, B**). Differences in the number of nodes and vessels per node were not significant (**Fig. 7C,D**).

**Figure 7.**
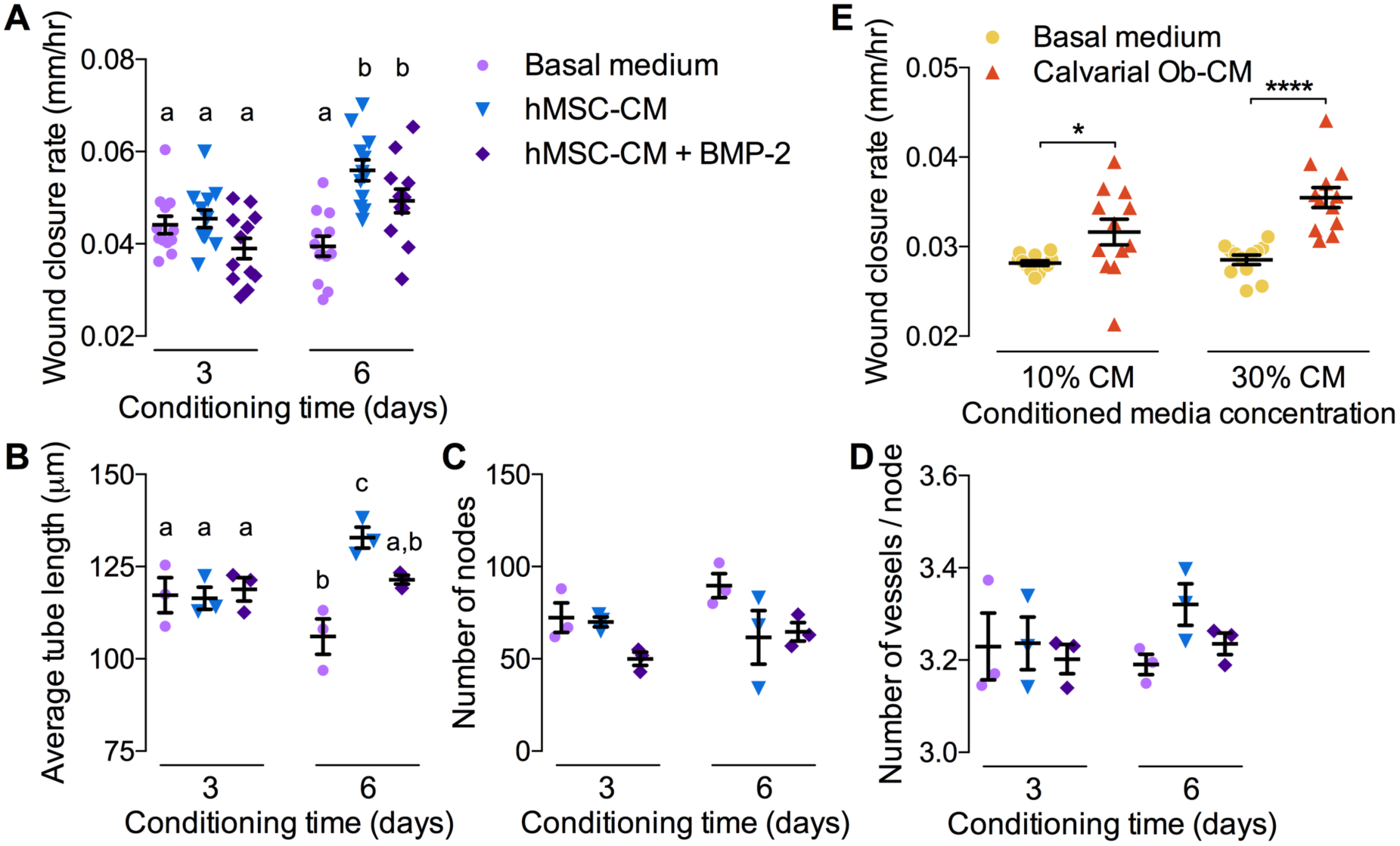
Effects of hMSC-and calvarial osteoblast-conditioned media on angiogenesis. Conditioned media was collected from hMSCs with or without 100 ng/ml BMP-2 (hMSC-CM + BMP-2 and hMSC-CM, respectively) after 3 or 6 days culture. Migration and tube formation were evaluated using a mix of conditioned media (30% by volume) and ECFC basal medium (BM). **A**) ECFC wound closure, quantified as wound closure rate (n=12/group in N=4 independent experiments). **B-D**) Tubular network formation (N = 3/group). **B**) Average tube length. **C**) Number of nodes. **D**) Number of vessels per node. Similarly, conditioned media was collected from mouse calvarial osteoblasts after 6 days culture. Migration and tube formation were evaluated using a mix of conditioned media (10% or 30% by volume) and ECFC basal medium (BM). **E**) ECFC wound closure, quantified as wound closure rate (n=12/group in N=4 independent experiments). Each replicate is shown with summary statistics represented as mean ± s.e.m. Significance indicator letters shared in common between groups indicate no significant difference by one-way ANOVA and Tukey’s multiple comparisons test.

To test whether conditioned media from mouse calvarial osteoblast-derived cells (COb) could similarly induce angiogenesis in a paracrine fashion, we cultured COb for 6 days and collected conditioned media, which was mixed to 10% or 30% by volume with ECFC basal medium. Wound migration and tube formation assays were then performed as above. Calvarial cell conditioned medium at 30% significantly increased wound closure rate (**Fig. 7E**), but did not alter tubular network formation (**Figure S1**).

### Effects of BMP-2 and/or ECFC delivery on vascularized bone formation at heterotopic sites

Together, these data suggest that BMP-2-induced neovascularization is a consequence of paracrine signaling from osteoprogenitor cells and osteoblasts rather than direct stimulation of endothelial cell angiogenesis. BMP-2 was first identified as an osteoinductive growth factor by its ability to induce bone formation at heterotopic sites, such as within the subcutaneous space.^23^ Similarly, the subcutaneous implant model is often used to study angiogenesis *in vivo*. For example, we recently showed that human ECFCs are capable of forming a functional human neovascular plexus that inosculates with the host vasculature when implanted into the subcutaneous flank of immunocompromised mice.^19^ Unlike orthotopic bone defects, in which endogenous osteoprogenitor cells and osteoblasts are relatively abundant, the subcutaneous model provides an opportunity to investigate osteogenic-angiogenic coupling in a niche that is relatively deficient in endogenous osteoprogenitor cells.

To this end, we developed collagen matrices, polymerized to fit into polysuflone disks (**Fig. 8A**), for subcutaneous implantation into the rear flank of RNU nude rats (**Fig. 8B**). The matrices were polymerized with or without ECFCs (384,000 cells/construct) and with or without BMP-2 (24 ng per construct) and cultured for 24 hours prior to subcutaneous transplantation for four weeks (N=6 per condition). Each of six animals received four implants, one from each group. At week 4, bone formation and neovascularization were evaluated by microCT angiography (**Fig. 8C**).

**Figure 8.**
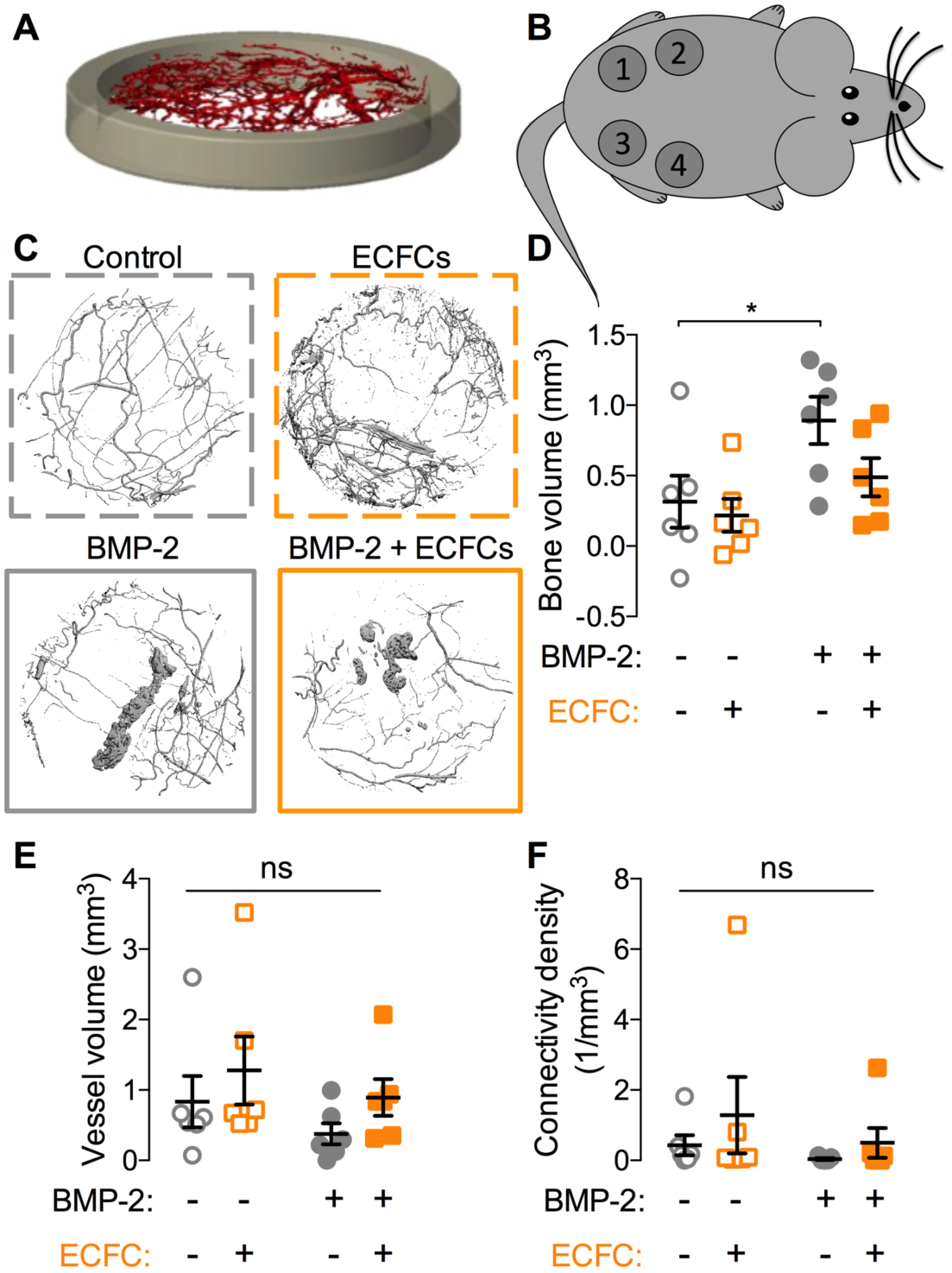
Effects of BMP-2 and/or ECFC delivery on heterotopic vascularized bone formation. A) Three-dimensional isometric rendering of the heterotopic chamber. B) Diagram of construct placement. Each of six animals received four implants, one from each group. Positioning of the impants was balanced to avoid bias by location. C) Representative microCT reconstructions of bone and blood vessel formation within implanted hydrogels. Dashed border indicates conditions without BMP-2 and solid borders indicate BMP-2 treated matrices. D) Neovascular volume. E) Connectivity density. Each replicate is shown with summary statistics represented as mean ± s.e.m. * indicates (p ≤ 0.05) by two-way ANOVA and Bonferroni multiple comparisons test. N=6.

BMP-2 delivery significantly induced heterotopic bone formation, but, as expected, ECFC delivery did not (**Fig. 8D**). Neither BMP-2 delivery, ECFC delivery, nor their combination significantly affected neovascular volume or connectivity measured by post-decalcified microCT angiography (**Fig. 8E,F**).

## Discussion

In this study, we evaluated the capacity of BMP-2 delivery to induce vascularized bone regeneration and tested whether the effects were primarily direct or indirect. We found that BMP-2 induced bone formation and neovascularization in large segmental bone defects. BMP-2 failed to induce endothelial cell migration or tube formation *in vitro*, but conditioned medium from either hMSCs or mouse calvarial osteoblast-derived cells significantly enhanced endothelial cell functions key to angiogenesis. Finally, we developed a new subcutaneous chamber model and showed that BMP-2 and/or ECFC delivery to a heterotopic site that is largely deficient in endogenous osteoprogenitor cells failed to induce neovascularization, in spite of BMP-2-induced osteogenesis.

Taken together, these observations identify challenges for endogenous bone tissue engineering approaches involving growth factor delivery alone. As observed previously,^24,25^ we found that BMP-2 robustly induces bone formation in supportive environments and concomitantly enhances neovascular growth. However, here we conclude that BMP-2-induced angiogenesis occurs primarily through paracrine mechansims secondary to osteoprogenitor activation rather than by direct stimulation of angiogenesis. Thus, in regenerative contexts featuring both adequate mesenchymal progenitor cell and adjacent vascular supply, BMP-2 may be sufficient to induce bone formation and physiologic neovascularization, but for injuries in which endogenous cell sources is compromised, including massive defects or multi-tissue polytrauma, BMP-2 delivery alone may be inadequate.

The influence of BMP-2 on angiogenesis is controversial. We and others have shown previously that BMP-2 delivery to rat segmental bone defects can significantly enhance neovascular network formation.^11,26^ However, other studies observed no effects of BMP-2 delivery on bone defect vascularization.^27,28^ Here, we tested the effects of BMP-2 on neovascularization in large bone defects *in vivo*, on isolated endothelial cells *in vitro*, and on ectopic angiogenesis in a niche poor in osteoprogenitor cell supply. Together, our results suggest that the primary role of BMP-2 in neovascularization during bone repair is through progenitor cell and osteoblast activation, which then stimulate angiogenesis in turn through paracrine signaling.

Studies on the direct effects of BMP-2 in endothelial cells are highly contradictory. Some report a pro-tubulogenesis response without enhancing motility,^16^ while others observed a stimulatory effect on migration and tube formation in human microvascular endothelial cells^29^ and human umbilical vein endothelial cells (HUVECs).^13^ BMP-2 has not been shown to effect HUVEC proliferation^13^ and only marginally increased DNA synthesis of HAEC and HUVECs.^16^ These results suggest that BMP-2 can serve as a chemoattractant and proangiogenic cytokine, but is not uniformly a mitogenic activator for these endothelial cell types. We did not detect effects of BMP-2 on ECFC migration or tubulogenesis, despite being responsive to BMP-2 at the level of signaling. Endothelial cells have BMP-2 receptors IA, IB, and II as well as the endothelial-specific TGFβ type III receptor (endoglin), which facilitates the binding of BMP-2 to its receptor.^30^ Our data confirm that ECFCs express functional BMP-2 receptors and that the BMP-2 used was functional and at a sufficient dose; however, BMP-2 did not induce a detectable angiogenic effect in ECFCs using these assays. In contrast, conditioned medium from either human mesenchymal stromal or mouse calvarial osteoblast cells enhanced ECFC migration. Conditioned medium from BMP-2-treated mesenchymal cells moderately, but not significantly, reduced measures of angiogenesis. As the data were not statistically significant, this effect could be incidental or could be associated with a shift from proliferation to differentiation status that could have reduced cell numbers compared to non-differentiated MSCs in growth medium.

Endothelial colony forming cells (ECFCs) are circulating endothelial cells that exhibit vasculogenic capabilities and home to sites of injury,^31,32^ motivating our use of this cell type in this study. In comparison to mature, quiescent vessel cell types including HAECs and HUVECs, ECFCs home from the bone marrow and incorporate into sites of vascular regeneration *in vivo*.^33,34^ These cells are also capable of forming blood vessels *de novo*.^19,35^ These observations motivated our selection of these cells as an *in vitro* model system to study wound-induced neovascularization.

Additionally, these cells have shown promise as a potential clinical therapeutic with their ability to incorporate into neovessels of regenerating tissues.^35^ ECFCs have been shown to provide paracrine support to mesenchymal stem cells prior to establishment of neovascularization^36^ and have been used in tissue-engineered scaffolds to create preformed vascular networks that have the ability to interconnect with the host vasculature *in vivo*.^37^ Further, coordinated spatial and temporal release of BMP-2 and VEGF enhanced osteogenesis and vasculogenesis of hMSCs and ECFCs within a patterned hydrogel implant, with the greatest response seen with VEGF released over 10 days and BMP-2 released over 21 days.^38^ This further supports the importance of the time at which angiogenic and osteogenic factors are presented. Our studies suggest that the angiogenic effects observed for ECFCs may not be primarily influenced by the presence of BMP-2.

Our data, including the paracrine signaling responses observed in MSC- and calvarial cell-conditioned media experiments, are consistent with recent studies showing that physiologic BMP-2 induces osteoblast expression of VEGF, and subsequent angiogenesis.^14^ Further vascular smooth muscle and endothelial cells have been shown to express BMP-2 during bone healing,^39^ forming a positive feedback interaction during fracture repair.^40^

### Ectopic bone formation and angiogenesis

Here, we used an extraosseous implantation site (subcutaneous chamber) to minimize the effects on neovascularization of the progenitor and osteoblast cells present in the defect model. While BMP-2 induced bone formation, we observed no detectable effect on endogenous or co-implanted vasculature, suggesting that neovascularization, independent of robust osteoprogenitor cell activation, was not significantly affected by the presence of BMP-2. Interestingly, the ability of rhBMP-2 to form bone in ectopic sites is inhibited by administration of TNP-470, an antiangiogenic agent that inhibits neovascularization.^41^ It is known that angiogenesis is critical for endochondral bone development;^42^ however, angiogenesis blockade, initiated after cell-recruitment and chondrogenesis but prior to osteogenesis, remained permissive of ectopic bone.^41^ Together, these data support the hypothesis that while angiogenesis is critical during early phases of endochondral bone healing, angiogenesis serves to recruit and support of BMP-receptor positive cells that would then be able to respond to locally delivered BMP-2.^41^

### Limitations

In these experiments, we evaluated neovascularization by microCT angiography, which is limited by voxel resolution (here, 15 μm). By threshold selection and partial volume effects we are likely able to detect vessels with diameters approaching this size, but this method is not able to resolve important microvessels, which may respond distinctly to BMP-2 treatment compared to vessels of larger size. We selected ECFCs as an endothelial cell model to mimic the cell types active during natural tissue regeneration. Other endothelial cell types, such as HMVEC or HUVEC may exhibit differential angiogenic responsiveness to BMP-2 treatment. While we have observed definitive evidence of translpanted ECFC inosculation into the mouse vasculature by lectin perfusion and immunostaining,^19^ microCT angiography may not have the resolution to resolve these neovessels. We selected a known healing dose of BMP-2 for the segmental defect studies,^43^ and BMP-2 concentrations *in vitro* that have been shown to robustly induce osteogenesis.^44,45^ However, the precise concentrations of BMP-2 experienced by invading neovessels during bone regeneration are unclear,^46,47^ and different BMP-2 concentrations may have direct angiogenic effects.

## Conclusions

Taken together, this study implicates paracrine signaling as the primary mechanism by which BMP-2 induces neovascularization during bone repair, and identifies endogenous neovascular and mesenchymal progenitor cell supply as important contextual considerations for the efficacy of BMP-2 delivery for bone regeneration.

## Supplementary Data

**Figure S1.**
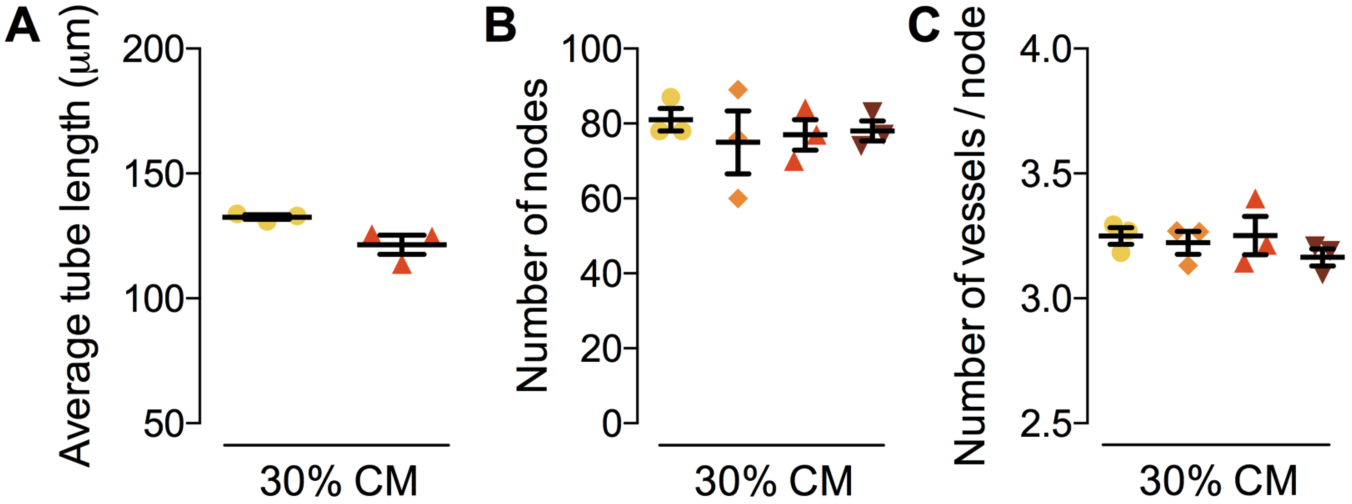
Effects of calvarial osteoblast-conditioned media on tubular network formation. Tube formation were evaluated using a mix of conditioned media (30% by volume) and ECFC basal medium (BM). **A-C**) Tubular network formation (N = 3/group). **A**) Average tube length. **B**) Number of nodes. **C**) Number of vessels per node.

